# MeSHHeading2vec: A new method for representing MeSH headings as feature vectors based on graph embedding algorithm

**DOI:** 10.1101/835637

**Authors:** Zhen-Hao Guo, Zhu-Hong You, Hai-Cheng Yi, Kai Zheng, Yan-Bin Wang

**Affiliations:** Xinjiang Technical Institute of Physics and Chemistry, Chinese Academy of Sciences, Urumqi 830011, China; University of Chinese Academy of Sciences, Beijing 100049, China; School of Computer Science and Technology, China University of Mining and Technology, Xuzhou, 221116, China; School of Cyber Science and Technology, Zhejiang University, Hangzhou 310000, Zhejiang

## Abstract

**Motivation:** Effectively representing the MeSH headings (terms) such as disease and drug as discriminative vectors could greatly improve the performance of downstream computational prediction models. However, these terms are often abstract and difficult to quantify.

**Results:** In this paper, we converted the MeSH tree structure into a relationship network and applied several graph embedding algorithms on it to represent these terms. Specifically, the relationship network consisting of nodes (MeSH headings) and edges (relationships) which can be constructed by the rule of tree num. Then, five graph embedding algorithms including DeepWalk (DW), LINE, SDNE, LAP and HOPE were implemented on the relationship network to represent MeSH headings as vectors. In order to evaluate the performance of the proposed method, we carried out the node classification and relationship prediction tasks. The experimental results show that the MeSH headings characterized by graph embedding algorithms can not only be treated as an independent carrier for representation, but also can be utilized as additional information to enhance the distinguishable ability of vectors. Thus, it can act as input and continue to play a significant role in any disease-, drug-, microbe- and etc.-related computational models. Besides, our method holds great hope to inspire relevant researchers to study the representation of terms in this network perspective.

## 1 Introduction

Technological advances over the past few decades, from high-throughput sequencing technologies to omics, have dramatically changed the paradigm of medicine and biology (Reuter, et al., 2015; Tyanova, et al., 2016). In particular, since the official launch of the Human Genome Project in the 1990s, the large-scale genomic, chemical and pathological data has brought novel insights for humans to re-recognize life processes (Collins, et al., 2003). However, the information overload caused by tremendous growth of data makes it difficult to take full use of existing knowledge and literature. For instance, a premier database called MEDLINE contains about 26 million records from more than 5,600 selected publications covering biomedical and life sciences to the present. So how to efficiently organize and manage the literature and explore the implicit value becomes a formidable challenge.

In response to this situation, the literature-based discovery (LBD) method was firstly proposed by Don R. Swanson which logically combines independent pieces of information to infer new interesting discoveries (Swanson, 1986). Many models were continuously developed to provide efficient and stable support for researchers such as co-occurrence-based approaches (Swanson and Smalheiser, 1997), semantic relation-based approaches (Hu, et al., 2010), Graph-based approaches (Cameron, et al., 2015) and Hybrid approaches (Torvik and Smalheiser, 2007).

For traditional LBD method, such as Medical Subject Headings (MeSH), Unified Medical Language System (UMLS) and etc. are often treated as auxiliary knowledge sources to improve the performance of the model. Despite significant progress has been made in this domain, most of them ignored the potential value behind the MeSH headings that itself is carefully designed. In addition, terms such as disease, drug and microbe are abstract entities that are difficult to be represented as concrete vectors as input for machine learning models. In this paper, we focus on analyzing MeSH to mine the hidden information. It is believed that this expert knowledge can be utilized to precisely quantify these terms.

MeSH is a kind of controlled and comprehensive vocabulary for subject indexing and searching books or journals in life sciences (Lipscomb, 2000). It was produced by National Library of Medicine (NLM) since 1960 and widely used around the world. More than half a century of heavy application has made MeSH increasingly perfect and made significant contributions to various fields. The MeSH consists of 3 parts including Main headings, Qualifiers and Supplementary Concepts. Main headings as the trunk of MeSH are used to describe the content or theme of the article. Qualifiers is the refinement of MeSH headings, i.e. how to be processed when it is in a specific area. Supplementary Concept is a complementary addition that is mostly related to drugs and chemistry. Some new substances have not yet become the main subject and will be included in Supplementary Concept to promote the integrity of MeSH. Here, we focus on discussing Main headings which consists of MeSH headings (descriptors), corresponding entry term and tree num.

MeSH Headings can be divided into 16 categories such as category A for anatomy, category B for organisms, category C for diseases, category D for Chemicals and Drugs, and etc. In MeSH tree structure, MeSH headings are organized as a “tree” with 16 top categories in which the higher hierarchy has the broader meaning and the lower hierarchy has the specific meaning. Compared with the tree structure, network (graph) is an more important data type which widely spreads in the real world and has been deeply researched (Cai, et al., 2018). Effective analysis of the Network not only can deeply understand the original graph, but also facilitate down-stream tasks such as node classification and relationship prediction. Hence, we construct the MeSH heading relationship network from tree structure through hierarchical tree num rules.

Graph embedding (network representation) is a kind of method to process the network problem which aims at transforming the node into low-dimensional vectors. In this process, it maximumly preserve both the local and global structure of the network. The mainstream Graph embedding algorithms can be roughly divided into 3 categories: factorization-based methods, random walk-based and deep learning-based methods (Goyal and Ferrara, 2018). The random walk-based graph embedding method is to use the random walk on the network to obtain a series of node paths to mimic the sentences or text. Then the Word2vec model can be applied to transform the node into vectors. The method of factorization takes the adjacency matrix as the structure of the graph, and obtains the node representation vectors by the method of matrix decomposition. Explosive research on deep learning has rapidly expanded its field to the network. The deep learning-based method is to carry out the feature capture and dimensional reduction tasks on node original representation to get the new low-dimensional vectors.

In this paper, the mainstream idea of using MeSH as a dictionary for indexing is abandoned, we transform the MeSH tree structure into a relationship network and implement 5 common graph embedding algorithms on it to represent the MeSH headings as vectors. In general, the whole process can be divided into 3 steps. Firstly, MeSH headings, tree num and entry terms were downloaded from National Library of Medicine (NLM) in September 22, 2019. Then we connected different Mesh headings through the rules of tree num to convert the tree structure to the relationship network. The label (category) of each node (Mesh heading) in the relationship network can be defined by the mode of its corresponding tree num. Secondly, the network has been briefly analyzed, including the number of nodes and edges, the distribution of node degrees and labels. Thirdly, we applied 5 network representation (graph embedding) algorithms including DeepWalk (Perozzi, et al., 2014), LINE (Tang, et al., 2015), SDNE (Wang, et al., 2016), LAP (Belkin and Niyogi, 2003) and HOPE (Ou, et al., 2016) to map the nodes into low-dimensional dense vectors which maximumly preserves the original network structure and the node relationship information. Then, we performed 2 types of tasks including node classification and relationship prediction. The node classification and relationship prediction tasks are used to assess the distinguishability of vectors between and within categories. In relationship prediction task, we performed drug-target interaction and miRNA-disease association prediction tasks to display that the term representation vectors can be as input for machine learning model. All results achieved by our method implied that the representation vectors generated by MeSH relationship network is efficient and reliable. High quality MeSH heading representation will definitely improve the prediction performance of existing computational models. At the same time, we hope that this work can provide novel insight to inspire relevant medical and life science researchers to mine the semantic information in MeSH through the network method. The flowchart is shown in the Fig. 1.

**Fig. 1.**
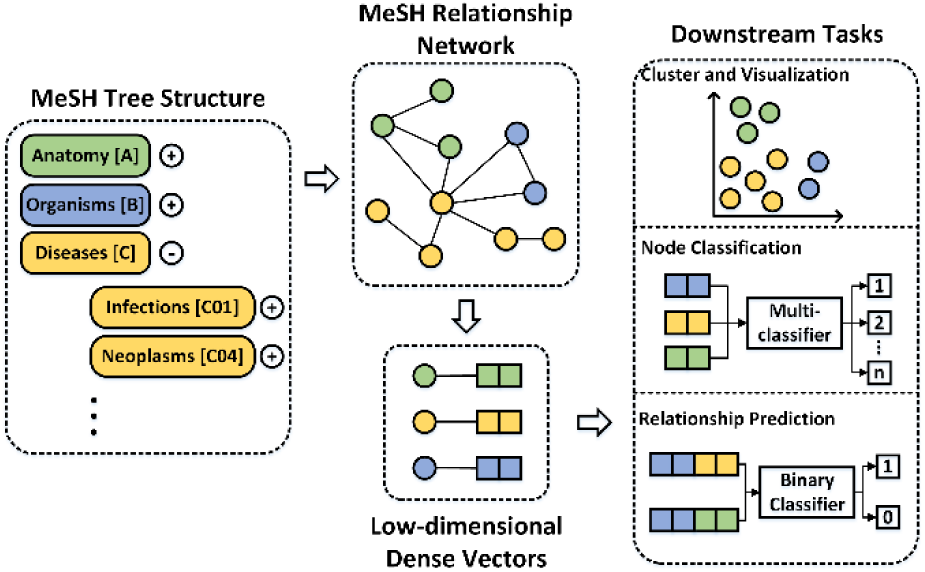
The flowchart of the proposed method includes three steps: construction, analysis and applications.

## 2 Materials & Methods

### 2.1 MeSH headings, tree numbers and entry terms

The Medical Subject Headings (MeSH) is a controlled and hierarchically-organized vocabulary directed by the National Library of Medicine (NLM) which is utilized for indexing, searching, and etc. in medical and life sciences. We downloaded MeSH headings, tree num and entry terms from NLM in September 22, 2019, and arranged them by routine standardized pretreatments including identifier unification and redundancy removal. After above operations, 29,349 Mesh Headings including their corresponding tree num and entry terms are congregated together for network construction.

Each MeSH heading can be descripted by one or more tree num to reflect its hierarchy in the tree structure and relationships with other MeSH headings. Tree num consists of letters and numbers, the first of which is uppercase letters represent category and the rest are made up of numbers. Each 3 digits represent a hierarchy in the tree structure. There are some MeSH headings such as lung cancer (C04.588.894.797.520, C08.381.540 and C08.785.520) are described by a single type of tree num, while others such as Reflex (E01.370.376.550.650, E01.370.600.550.650, F02.830.702 and G11.561.731) can be represented by different kinds of tree num. The labels can be labelled by the mode of the node’s tree num, Reflex will be given a label E.

Whenever the last hierarchy of tree num is removed, a new tree num and corresponding MeSH heading can be generated and contacted. The details can be seen the figure 2. Through the formation of this kind of relationship, a MeSH heading network consisting of 29,349 nodes and 39,784 edges can be constructed. For the sake of simplicity, we treat the mode of the tree num category of MeSH heading as its label.

**Fig. 2.**
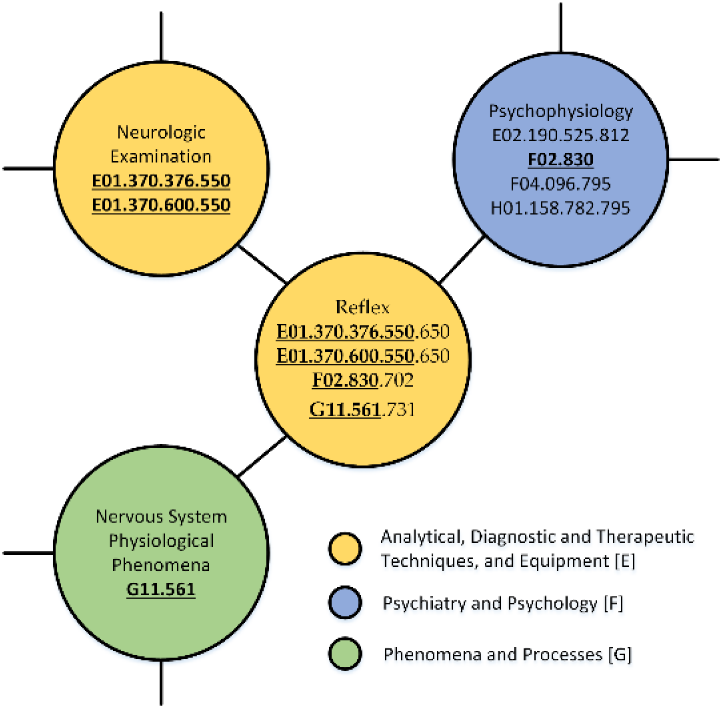
The construction of the MeSH relationship network. Reflex has 4 tree num including E01.370.376.550.650, E01.370.600.550.650, F02.830.702 and G11.561.731. The Neurologic Examination (E01.370.376.550, E01.370.600.550) can be obtained when the last 3 digits (.650 and .650) of Reflex (E01.370.376.550.650, E01.370.600.550.650) are removed. The category (label) of each MeSH heading is the mode of its corresponding tree num.

Entry term is a kind of synonym or similar vocabulary for MeSH headings. In order to unify identifiers and eliminate ambiguity, we create a MeSH Heading Term Correspondence Table to convert the entry terms to standard MeSH headings.

### 2.2 Benchmark datasets

2 benchmark datasets were collected as the benchmark datasets to be utilized to verify the performance of the proposed method.

#### 2.2.1 known drug-target interactions

28,211 known drug-target interactions were downloaded from DrugBank in May 8, 2019 (Wishart, et al., 2017). After standardizing the identifiers via the Correspondence Table and STRING database, we got of 7,739 different drugs and 4,975 different proteins. In order to avoid sparsity of associations, we selected drugs and proteins that are associated with more than 5 corresponding objects similar to the article described by Zhang *et al.* (Zhang, et al., 2018). Finally, we obtained 7,318 experimental valid drug-target interactions containing 641 different drugs and 317 different proteins.

#### 2.2.2 known miRNA-disease associations

35,547 known human miRNA-disease associations which consist of 1206 different miRNAs and 894 different diseases were downloaded from HMDD in May 8, 2019 (Huang, et al., 2018). Considering that the name of disease and miRNA in the original database are nonstandard such as “breast neoplasms” and “carcinoma, breast” are the same type of disease. After standardizing the identifiers via miRBase and the Correspondence Table described above, we obtained 11,109 experimental valid miRNA-disease associations containing 843 different miRNAs and 531 different diseases.

#### 2.2.3 Positive and negative samples

The experimental-validated interaction or association pairs are regarded as the positive samples and the randomly selected equal unlabeled pairs are treated as negative samples. This is a typical strategy that equalizes training samples and is widely used in bioinformatics (Ben-Hur and Noble, 2005). Each positive and negative sample is given a label 1 and 0, respectively.

### 2.3 Baseline methods

3 baseline methods including Morgan molecular fingerprint, k-mer and disease similarity are chosen to represent drug, protein and disease as vectors, respectively.

#### 2.3.1 Molecular Fingerprint

Molecular Fingerprint is one of the most popular methods to represent drugs by describing the structure of compounds. The basic idea is to segment the drug molecule and obtain structure fragments one after another. Then, these substructures are encoded into numbers according to certain rules, which can correspond to each of the binary strings. The whole binary string is used as the characterization of drug molecular structure. In this paper, fingerprint method is chosen as the baseline to represent the drug. The SMILES of each drug were downloaded from DrugBank and transformed into fingerprints by python package called RDKit (Landrum, 2013).

#### 2.3.2 K-mer method

For a long time, how to transform sequences efficiently and reliably into numerical representations is a formidable challenge. In this article, a widely used bassline method called k-mer is applied and the details of the algorithm are shown as follows.

For protein and miRNA, the sequences of them were downloaded from STRING (Szklarczyk, et al., 2018) and miRBase (Kozomara, et al., 2018), respectively. Inspired by Shen *et al.* (Shen, et al., 2007), we represent proteins and miRNAs as vectors by analyzing and normalizing their components. For proteins, we classified 20 amino acids into 4 groups according to the polarity of the side chain, including (Ala, Val, Leu, Ile, Met, Phe, Trp, Pro), (Gly, Ser, Thr, Cys, Asn, Gln, Tyr), (Arg, Lys, His) and (Asp, Glu). For miRNA, there naturally exist 4 types of nucleotides including Adenine (A), Cytosine (C), Guanine (G) and Uracil (U) in the sequence. Then, each miRNA or protein can be abstracted into a vector by the method k-mer, in which all dimensions represent the full permutation of k nucleotide combinations and the value of each dimension is the normalized frequency of the corresponding k-mer appearing in the sequence. Here, we set k to 3 and the dimension of the representation vector is 64 (4^3^).

#### 2.3.3 Disease similarity

Disease is an abnormal life activity process that occurs when a living organism is destructively affected by a certain cause. The semantic similarity of disease is a common method of abstracting disease into vectors (Guo, et al., 2019). For each disease, a Directed Acyclic Graph (DAG) can be constructed by the MeSH heading relationship in Section 2.1. Specifically, disease *D*’s ancestor nodes can be obtained by continuously removing the last hierarchy of its tree num. *D* and its ancestor nodes together constitute a DAG. Then the similarity between 2 diseases can be calculated according to the generalized Jaccard formula, *i.e.*, the larger the intersection, the more similar it is. According to the previous literature (Wang, et al., 2010), the specific calculation process is as follows:

For disease *D, DAG(D)* = (*D, N(D), E(D)*), *N(D)* is the point set that includes all *DAG(D)*’s diseases. *E(D)* is the edge set that includes all *DAG(D)*’s relationships. The semantic value contribution of disease d in the set *N(D)* to disease *D* can be defined as:

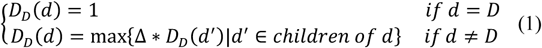

where Δ denotes a decline factor. In the *DAG(D), D* can be seen as the disease that contributes the most to its own semantic value and equals to 1, and the remaining diseases will contribute less and less to disease *D* as the distance increases. Then, the sum of the contributions of diseases which are in the set *N(D)* to *D* can be calculated as follows:

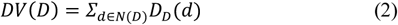

Finally, the similarity between diseases *m* and *n* can be calculated by the following formula:

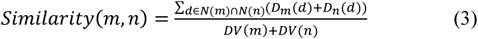

The disease similarity matrix of *k* rows and *k* columns containing *k* different diseases can be constructed, and the *i*-th row can be regarded as a representation vector of the *i*-th disease.

#### 2.3.4 Autoencoder

In order to unify the dimensions of the vector and obtain a higher quality representation, autoencoder is applied to map the drug fingerprint and disease similarity from original space to the low-dimensional space. Hidden layer representation *h* and output layer representation *y* can be calculated by the following formula:

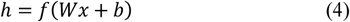

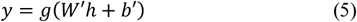

Where *x* is input, *W* and *b* are weights and thresholds, respectively, *f* and *g* are the activation functions. Loss function can be obtained by minimizing the error between input and output:

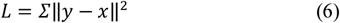

Finally, all drug fingerprint and disease similarity can be normalized to 64-dimensional vectors.

### 2.4 Graph embedding methods

Mesh heading relationship network is a complex heterogeneous network. Analysis of network can better help us understand this kind of unstructured data and benefit the exploration of the underlying knowledge. Graph embedding is an effective method to provide new insights on how to make good use the hidden information behind the graph. In this chapter we first give a graph embedding formal definition, and then briefly introduce several algorithms used in this paper.

A graph *G*(*V, E*) is a collection of vertices (node) set *V* = {*v*_1_, …, *v*_*n*_} and edge set 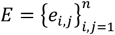. The aim of graph embedding is to find a mapping function *f*: *v*_*i*_ → *x*_*i*_ ∈ *R*^*d*^, where *d* ≪ |*V*|, and *X*_*i*_ = {*x*_1_, *x*_2_, …, *x*_*d*_} is the embedded vector that captures the structural of vertex *v*_*i*_

In this paper, we apply 5 kinds of graph embedded methods on the network to perform downstream tasks including node classification and relationship prediction.

DeepWalk (DW) obtains a series of node sequences through random walks of vertexes in the network and inspired by the Skip-Gram model to analogize these paths to sentences for representation learning. The goal is to learn a latent representation and the mapping function is:

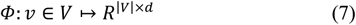

The problem then, is to estimate the likelihood:

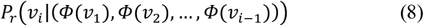

The recent relaxation in language modeling turns the prediction problem and this yields the optimization problem:

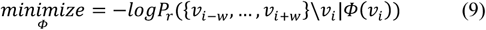

Large-scale Information Network Embedding (LINE) is an efficient network representation learning algorithm that is quite different from random walk-based method. Low-dimensional dense vectors can be obtained by LINE by preserving first-order and second-order proximity. For first-order, the objective function can be defined as follows:

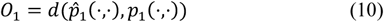

For the edge *e*_*i,j*_ which from vertex *v*_*i*_ to vertex 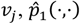 and *p*_1_(·,·) are the empirical and joint distribution respectively between the latent embeddings 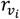 and 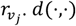 is the distance between above 2 distributions.

For second-order, the objective function can be defined as follows:

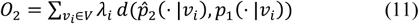

where 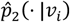 and *p*_1_(· |*v*_*i*_) are empirical and context conditional distribution for each *v*_*i*_ ∈ *V* under the model by vertex embeddings. For the sake of simplicity, λ_*i*_ is set to the degree of the vertex *i*.

Structural Deep Network Embedding (SDNE) is a semi-supervised deep autoencoder consisting of supervised and unsupervised component that can capture the nonlinear structure from the network. For the supervised part, the objective function can be defined as follows:

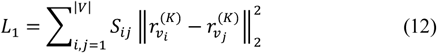

Where 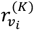 is the K-th layer representation of *v*_*i*_.

For the unsupervised part, the objective function can be defined as follows:

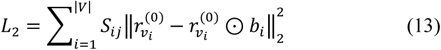

where 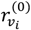 is the representation of *v*_*i*_ and *b*_*i*_ is a weight vector.

Finally, the joint objective function can be defined as follows:

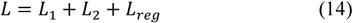

where *L*_*reg*_ is a regularization term to prevent overfitting.

High-order Proximity Preserved Embedding (HOPE) captures high order proximity of asymmetric transitivity in direct graph and symmetric transitivity in undirect graph. To achieve this goal, HOPE can obtain 2 vertex representation vectors *U*^*s*^, *U*^*t*^ ∈ *R*^|*V*|×*d*^, where *U*^*s*^ and *U*^*t*^ are called source and target vectors. The objective function can be defined as follows:

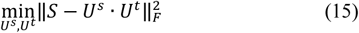

The structure of the reserved graph can be considered as the similarity of the reserved nodes. Laplacian Eigenmaps is an embedding algorithm which obtain the representation vector when the similarity parameter *W*_*ij*_ is high. The objective function can be defined as follows:

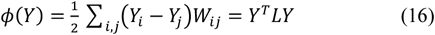

## 3 Results

### 3.1 Evaluation criteria

The MeSH relationship network consisting of nodes and the edges contains a wealth of medical and biological knowledge. After mining the content by embedding algorithms, low-dimensional dense representation vectors can be used for downstream tasks such as visualization, node classification and relationship prediction. How to evaluate the merits and demerits of the proposed method in a fair and comprehensive way becomes a formidable challenge.

Firstly, we briefly analyzed the MeSH relationship network. Secondly, we not only perform the node classification in the whole network, but also extract drug and disease representation vectors to carry out the relationship prediction tasks. Both of them aim at evaluating the distinguishability of vectors. High-quality representation vectors make it easier to construct the classifier to make prediction results more accurate. The details of the results can be seen in the following section.

Meanwhile, we applied a wide range of evaluation criteria to effectively assess the performance of our method (Wang, et al., 2019). Cross validation is a widely used method to measure model ability (Guo, et al., 2019; You, et al., 2017). For 5-fold cross-validation, the whole dataset is divided into 5 mutually exclusive subsets of roughly size, each subset is treated as the test set for evaluation in turn, the others are treated as the training sets for the model construction. At the same time, we draw ROC (receiver operating characteristic curve) and PR (Precision - Recall) to calculate AUC (area under ROC) and AUPR (area under PR) respectively in order to visualize experimental results and facilitate comparison with other methods. In addition, a wide range of evaluation criteria including accuracy (Acc.), sensitivity (Sen.), specificity (Spec.), precision (Prec.) and MCC have been adopted to evaluate our approach more generally.

### 3.2 Network analysis

The MeSH heading relationship network is a heterogeneous network consisting of 29,349 nodes and 39,784 edges, where the nodes are included by 16 different kinds of descriptors. Node degree refers to the number of edges associated with the node, also known as correlation degree. The occurrence number of the node and degree can be statistics and visualized as the fig. 3. In short, it fits the long tail distribution.

**Fig. 3.**
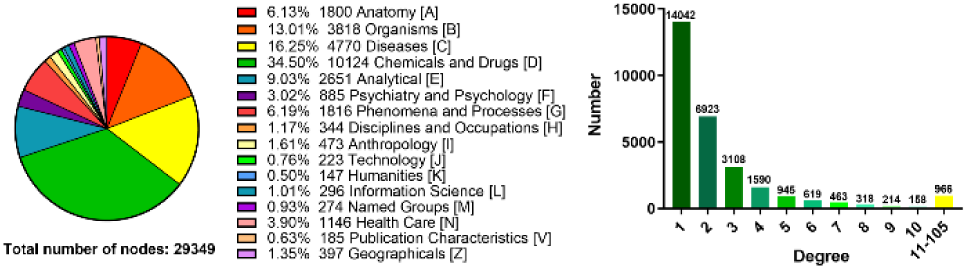
Distribution of node type and node degree in the relationship network.

### 3.2 Application 1: MeSH headings classification

As mentioned above, each node (MeSH heading) can be represented as a low-dimensional dense vector by graph embedding algorithm and can be labelled by the mode of its tree num. We want to verify the pros and cons of different graph embedding algorithms through the node classification experiment.

Specifically, 5 graph embedding algorithms including DeepWalk, LINE, SDAE, LAP and HOPE are applied on the relationship network to represent the nodes as 64-dimensonal vectors. Then, 80% of the nodes and the corresponding labels are utilized to construct the multi-classifier, and the remaining 20% of the nodes and the corresponding labels are used for testing. Although there exist some noises and errors in labels, the accuracy of the classifier can reflect the quality of the representation vectors to some extent. The results including ACC and LOSS are shown in the following figure 4.

**Fig. 4.**
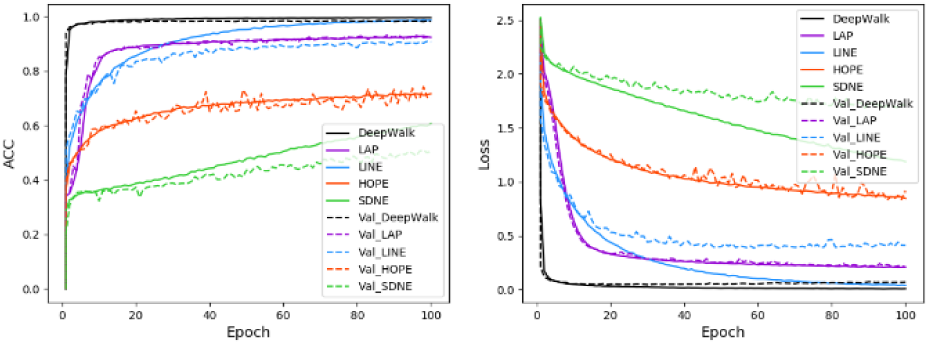
The training and valid performance of node classification task achieved by different graph embedding methods.

The node classification task reflects the distinguishability between different type of term representation, such as anatomy and organisms. Compared with other method, DeepWalk obviously achieved the most competitive performance which demonstrate that DeepWalk can indeed capture global structure and differences between various labels in the whole network.

The keras library was applied to construct this multi-classifier. An artificial neural network with 2 layers was built where each layer consists 512 neurons. The parameters including loss, optimizer, batch size and epochs are set to categorical crossentropy, RMSprop, 1024 and 100, respectively.

### 3.3 Application 2: representation for relationship prediction

#### 3.3.1 Drug representation for drug-target interaction prediction

In this section, we choose drug-target interaction prediction as a specific research subject to evaluate the quality of the drug representation vector. Specifically, each drug and protein can be represented as a 64-dimenonal vector by graph embedding and k-mer method. We also treated drug Morgan molecular fingerprint (FP) method as a baseline for comparison. Then, each drug-target interaction pair is a 128-dimensional vector by concatenating drug and target. 5-fold cross validation was applied to evaluate the performance of each method. Random Forest is chosen as the classifier to carry out the interaction prediction task. The details of the results can be seen in the following table 2 and fig. 5.

**Table 1.** The test performance of different graph embedding methods on the node classification task.

| Test Performance | SDNE | HOPE | LINE | LAP | DW |
| --- | --- | --- | --- | --- | --- |
| ACC | 0.5056 | 0.7003 | 0.9068 | 0.9284 | **0.9824** |
| Loss | 1.7108 | 0.9164 | 0.4130 | 0.2105 | **0.0722** |

**Table 2.** The performance of different graph embedding methods under 5-fold cross validation on the drug-target interaction prediction.

| Method | Acc. (%) | Sen. (%) | Spec. (%) | Prec. (%) | MCC (%) |
| --- | --- | --- | --- | --- | --- |
| FP | 77.49±0.32 | 72.45±0.85 | 82.53±0.59 | 80.58±0.43 | 55.27±0.60 |
| HOPE | 76.33±0.84 | 72.63±1.29 | 80.04±1.02 | 78.44±0.94 | 52.82±1.69 |
| LAP | 77.78±1.00 | 73.07±1.18 | 82.48±1.10 | 80.66±1.13 | 55.80±2.00 |
| LINE | 76.08±0.46 | 69.88±1.05 | 82.29±0.80 | 79.79±0.63 | 52.59±0.90 |
| SDNE | 75.63±0.41 | 70.37±0.57 | 80.88±0.84 | 78.64±0.70 | 51.54±0.86 |
| **DW** | **78.61±0.55** | **73.24±0.87** | **83.97±1.02** | **82.06±0.89** | **57.55±1.13** |

**Fig. 5.**
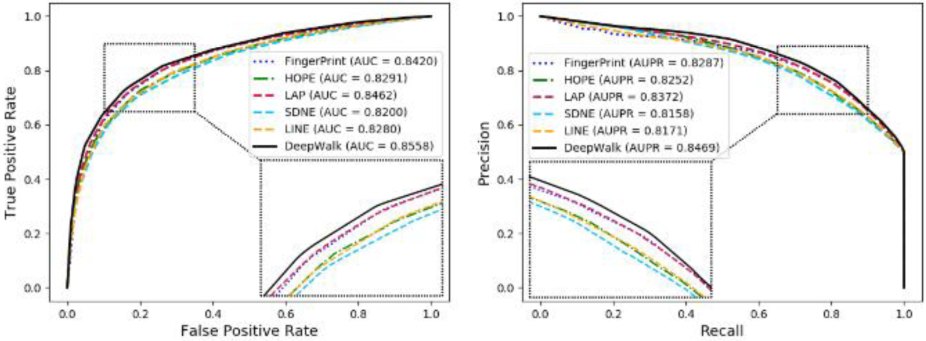
ROCs, AUCs, PRs and AUPRs of drug-target interaction prediction achieved by different graph embedding and drug Morgan molecular fingerprint methods.

Compared with the node classification, the relationship prediction task reflects the distinguishability between the same type of term representation. In general, DeepWalk and LAP achieved more competitive prediction effects. Considering the traditional method of analyzing the chemical structure of drugs, the satisfactory results proves that the proposed representation is novel and can adequate characterize the drug by semantic. We believe it will open up a new paradigm for semantic representation of drugs.

#### 3.3.2 Disease representation for miRNA-disease association prediction

In this section, we choose miRNA-disease association prediction as a specific research subject to evaluate the quality of the disease representation vector. Specifically, each miRNA and disease can be represented as a 64-dimenonal vector by k-mer and graph embedding method. We also performed disease similarity (DS) method as a baseline for comparison. Then, each miRNA-disease association pair is a 128-dimensional vector by concatenating miRNA and disease. 5-fold cross validation was chosen to evaluate the performance. Random Forest is applied as the classifier to carry out the association prediction task. The detail results of all methods can be seen in the following table 3 and fig. 6.

**Table 3.** The performance of different graph embedding methods under 5-fold cross validation on the miRNA-disease association prediction.

| Method | Acc. (%) | Sen. (%) | Spec. (%) | Prec. (%) | MCC (%) |
| --- | --- | --- | --- | --- | --- |
| DS | 82.42±0.28 | 79.01±0.75 | 85.84±0.92 | 84.81±0.74 | 65.01±0.59 |
| HOPE | 78.12±0.82 | 76.13±0.96 | 80.11±1.34 | 79.29±1.13 | 56.28±1.66 |
| LAP | 81.53±0.46 | 78.07±0.49 | 84.99±0.61 | 83.88±0.59 | 63.22±0.93 |
| LINE | 81.89±0.46 | 78.69±0.99 | 85.07±0.96 | 84.07±0.78 | 63.91±0.93 |
| SDNE | 82.68±0.62 | 79.17±1.61 | 86.18±1.42 | 85.16±1.16 | 65.53±1.22 |
| DW | **83.02±0.53** | **79.95±0.24** | **86.09±0.97** | **85.19±0.88** | **66.17±1.09** |

**Table 4.** The performance of different feature under 5-fold cross validation on the miRNA-disease association prediction.

| Method | Acc. (%) | Sen. (%) | Spec. (%) | Prec. (%) | MCC (%) |
| --- | --- | --- | --- | --- | --- |
| Attribute | 83.19±0.48 | 79.75±1.05 | 86.63±0.36 | 85.65±0.30 | 66.55±0.91 |
| Behavior | 83.56±0.82 | 77.23±1.30 | 89.89±0.92 | 88.43±0.97 | 67.67±1.62 |
| Both | **83.98±0.59** | **78.57±1.63** | **89.39±0.73** | **88.11±0.59** | **68.38±1.05** |

**Fig. 6.**
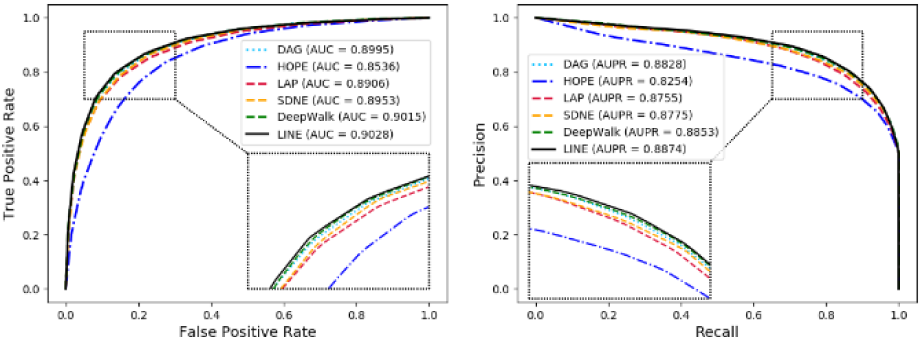
ROCs, AUCs, PRs and AUPRs of miRNA-disease associations prediction achieved by different graph embedding and DAG methods.

**Fig. 7.**
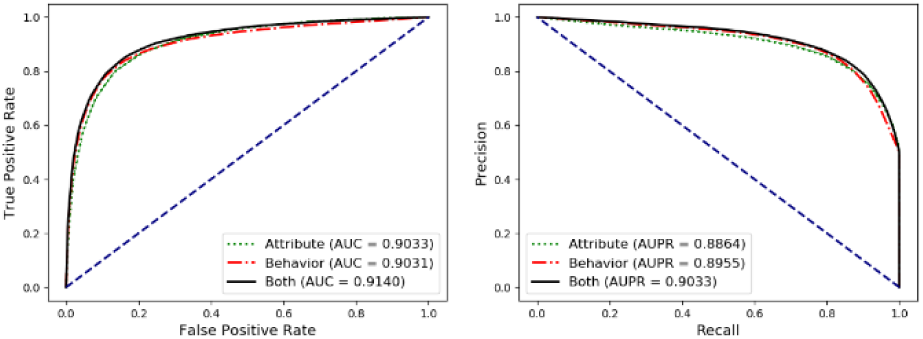
ROCs, AUCs, PRs and AUPRs achieved by different feature.

Briefly, the LINE method achieved the most remarkable with average Acc., Sen., Spec., Prec. and MCC of 83.02, 79.95, 86.09, 85.19, 66.17 and 90.28. The corresponding standard deviations of above evaluation criteria are 0.53, 0.24, 0.97, 0.88 and 1.09. The brilliant performance of the proposed method indicate that the representation vector generated by MeSH relationship network can be used as an independent carrier to characterize disease. Meanwhile, the lower standard deviation implied that the novel model was robust and stable.

Despite the performance improvement relative to the disease similarity is weak, the disease graph embedding representation has 3 obvious advantages. Firstly, compared with the similarity-based method, the graph-based method has faster calculation speed and less resource occupation. Secondly, the similarity-based method needs to be manually adjusted the decay factor while graph-based method needs less parameters to be set. Thirdly, graph-based method needs to be recalculated when facing a new sample, but the graph-based method can be generated once for permanent use.

#### 3.3.3 As addition information to enhance the ability of disease representation

In this section, we choose miRNA-disease association prediction as a specific research subject to prove that our representation method of disease can be utilized as the additional information. Specifically, inspired by Guo *et al.* (Guo, et al., 2019), each miRNA and disease can be represented by 2 kinds information including the behavior and the attribute feature. The behavior feature is the main information that is proposed by the idea of collaborative filtering or similarity theory. It is known that miRNAs with similar functions are often associated with similar diseases and vice versa. Then each miRNA and disease can be represented as a 64-dimensonal vector by known miRNA-disease associations through LINE method. The attribute feature is the additional information including the RNA sequence, disease semantics, drug chemical structure and etc. The attribute feature of each node can be represented as a 64-dimensonal vector by miRNA sequence learned by k-mer and disease semantics learned by LINE. Then each miRNA and disease can be viewed as a 128-dimensonal vector by connecting behavior and attribute feature. Finally, each miRNA-disease association pair is a 256-dimensional vector by concatenating miRNA and disease. 5-fold cross validation was applied to evaluate the proposed method. Random Forest classifier is chosen to carry out the association prediction task. The details of the results can be seen in the following figure and table.

The results demonstrated that the attribute feature (disease semantics graph embedding representation) can play an auxiliary role. The representation vector combined the two feature is easier to distinguish, which can improve the prediction performance of the computational model.

## Conclusion

Obtaining distinguishable vectors as the input of the computational prediction model has always been a hot topic of concern. Existing methods which manually define and measure similarity are limited considering the time and space complexity. In this paper, we constructed a MeSH heading relationship network and implemented 5 kinds of graph embedding algorithms on it. Then the qualities of the vectors were evaluated based on the relationship network itself and the 2 benchmark datasets including drug-target interaction and miRNA-disease association. Obviously, the results of relationship prediction prove that the semantic representation of terms such as disease can not only be used as independent carrier for input, but also as additional information to enhance the distinguishability of vectors. Despite the limited performance of the upgrade, compared with the previous feature generation approach such as similarity-based or chemical structure method, the proposed method is a fully automatic and pure semantic way, which will bring new enlightenment to relevant researchers. Predictably, MeSHHeading2vec can be viewed as a foundation to establish interesting connections between network and semantic in both computer and life sciences. Briefly, our method will establish valuable insights in MeSH heading representation and disease-, drug- and etc.-related computational prediction model, bring beneficial inspiration to relevant scholars and expand the computational omics research paradigm.

## Acknowledgements

We thank anonymous reviewers for very valuable suggestions.

